# A New Type of Nonsuppressible Viremia Produced by HIV-Infected Macrophage

**DOI:** 10.1101/2025.09.02.673877

**Authors:** Matthew J Moeser, Olivia D Council, Nathan Long, Laura Kincer, Ann M. Dennis, Joseph Eron, David Wohl, Claire E. Farel, Julie Nelson, Abbas Mohammadi, Behzad Etemad, Jonathan Z. Li, Hugh McGann, Erasmus Smit, Meredith Clement, Tat Yau, Prema Menezes, Natalie M. Bowman, Shuntai Zhou, Sarah B. Joseph

## Abstract

**Background:** HIV-1 RNA typically declines rapidly after initiation of antiretroviral therapy (ART); often reaching undetectable levels within a few weeks and remaining undetectable by standard assays. However, some patients on ART have persistent nonsuppressible viremia (NSV) that does not respond to treatment optimization or intensification. NSV can emerge at the time of ART initiation (*primary NSV*) or after being ART-suppressed (*secondary NSV*). Here, we examine mechanisms producing *primary NSV* in four people on ART.

**Methods:** Blood samples were collected from four participants who, despite being adherent to ART, required approximately a year or more to become virologically suppressed. Viral RNA and proviral DNA genomes were sequenced to examine HIV-1 drug resistance, genome intactness and genetic diversity. The ability of HIV-1 Envs to facilitate efficient entry into cells expressing low levels of CD4 (a proxy for macrophage tropism) was assessed.

**Results:** Before ART, the blood contained HIV-1 RNA genomes that were adapted to replication in CD4+ T cells and rapidly decayed after ART initiation. During ART, the blood contained HIV-1 genomes that were drug sensitive, genetically diverse, macrophage-tropic, not evolving and often had defects in *vpr*.

**Conclusions:** Our results suggest that in individuals with *primary NSV*, ART stopped virus replication, but large pools of long-lived, HIV-infected macrophage continued to produce virus. This is mechanistically distinct from *secondary NSV* produced by CD4+ T cell clones. In addition, defects in *vpr* independently accumulation in macrophage-tropic lineages found in three participants, suggesting that *vpr* may impact survival of, or virus production from, HIV-infected macrophage.

**SUMMARY:** People with HIV-1 on ART can experience persistent, nonsuppressible viremia (NSV). We identified four people with a previously uncharacterized form of NSV that emerged at the time of ART initiation (primary NSV) and is notable for its genetic diversity, drug sensitivity, and macrophage tropism. This is mechanistically distinct from previously described cases of NSV produced by CD4+ T cells in people who were previously ART-suppressed (secondary NSV).

## INTRODUCTION

Antiretroviral therapy (ART) is highly effective at suppressing HIV-1 RNA below the limit of quantification (LOQ) of standard clinical tests (∼20 to 50 HIV-1 RNA copies/mL [cp/mL]). However, some people with HIV-1 (PWH) are unable to quickly suppress HIV-1 RNA and others fail to maintain virologic suppression during ART [1]. Nonadherence or the presence of transmitted [2] or acquired [3] drug resistance mutations (DRMs) can prevent PWH from suppressing or maintaining suppression of HIV-1 RNA during ART. However, some people adherent to ART have persistent HIV-1 RNA in their plasma that is not drug resistant and cannot be suppressed by ART optimization or intensification; a condition termed nonsuppressible viremia (NSV) [4–7]. NSV can emerge at the time of ART initiation (*primary NSV*) or after an individual was ART-suppressed (*secondary NSV*). *Secondary NSV* has been extensively studied in recent years and is characterized by low to moderate levels of HIV-1 RNA in plasma (typically 200 to 1000 RNA cp/mL) [4–7] and can be composed of replication competent [4, 6] or defective viruses [7]. Analyses of HIV-1 integration sites in circulating CD4+ T cells suggest that *Secondary NSV* is often produced by clonally expanded CD4+ T cells [4, 5, 7].

In this study we describe four participants with a *primary NSV* that surprisingly was produced by virus production from HIV-infected macrophage. After starting ART, each participant had HIV-1 RNA levels that either declined slowly or did not decline despite having drug sensitive virus and well documented adherence. In contrast to previously described cases of *secondary NSV,* which were produced by virus expression from CD4+ T cell clones, the participants in this study with *primary NSV* had virus populations in their plasma that were genetically diverse, distinct from virus produced by circulating CD4+ T cells and adapted to replication in macrophage (M-tropic).

## METHODS

### Sample Collection

Samples were collected as part of clinical care at three sites (Table 1). Remnant samples were then deidentified for use in this study. Plasma samples were stored at -80 °C and PBMCs were isolated by Ficoll purification and stored at ≤ -150 °C. Viral load testing was performed at each site using standard clinical assays. Participants (P1, P2, P3 and P4) gave informed consent under protocols approved by the local institutional review boards at each site. P1, P3 and P4 were referred by physicians to assess drug resistance and the source of persistent viremia. P2 was enrolled in the Tropism of HIV-1 Inflammation and NeuroCognition (THINC) study and previously described in a study of CSF compartmentalization [8]. Participants from the University of North Carolina (UNC) were enrolled as part of the UNC Center For AIDS Research Clinical Cohort. P1 received intermittent ART for more than 10 years before initiating ART in this study (Supplemental Table 1).

**Table 1:** Clinical Characteristics of Participants.

|  | **Participant 1 (P1)** | **Participant 2 (P2)** | **Participant 3 (P3)** | **Participant 4 (P4)\*** |
| --- | --- | --- | --- | --- |
| **Site** | UNC Hospitals,<br>Chapel Hill, USA | UNC Hospitals,<br>Chapel Hill, USA | LSU Health Sciences<br>Center New Orleans, USA | Waikato Hospital<br>Hamilton, NZ |
| **Sex at Birth** | Male | Male | Male | Male |
| **Year Enrolled** | 2021 | 2011 | 2022 | 2023 |
| **HIV-1 Subtype** | B | B | B | B |
| **ART Regimen** | TAF/FTC/DTG/DRV/<br>COBI | TDF/FTC/DRV/r | BIC/FTC/TAF | FTC/TDF/DTG |
|  |  |  | FTC/TDF/DTG | FTC/TDF/DTG/DRV/r |
|  |  |  | CAB/RPVFTC/TAF | AZT/CAB/RPV/LEN/DRV/r<br>/IBA |
|  |  |  | CAB/RPV/DRV/COBI/FTC<br>/TAF |  |
|  |  |  | DRV/RTV/DTG/3TC |  |
| **OIs/Diseases**<br>**Associated With**<br>**Immunosuppression**<br>**and Other**<br>**Conditions** | EBV DNA+ CSF,<br>Transverse Myelitis | None | dMAC<br>KS<br>PJP<br>Otitis externa | dMAC<br>EBV adrenal tumor<br>Candidiasis<br>Microsporidiosis<br>CMV Retinitis |
| **At ART Start** |  |  |  |  |
| **Age (Years)** | 47 | 40 | 31 | 29 |
| **Viral load**<br>**(RNA cp/mL)** | 1,386,132 | 821,019 | 858,000 | 2,100,000 |
| **CD4 T cell**<br>**count (cells/uL)** | 90 | 10 | 16 | 12 |
| **6 months post ART** |  |  |  |  |
| **Viral Load**<br>**(RNA cp/mL)** | 1,562 | 710 | 13,300 | 14,533,087 |
| **CD4 T cell**<br>**count (cells/uL)** | 232 | 186 | 139 | 22 |
| **Time to Viral**<br>**Suppression**<br>**(weeks)** | 50 | 44 | 144 | Not Suppressed<br>by 57 weeks |

\*Participant 4 passed away after the end of the study period.

### HIV-1 Genetic Diversity and Bioinformatic Analyses

Diversity and drug resistance were assessed using deep sequencing with Primer ID Miseq [9]. Briefly, pools of primers with unique 11 base tag (Primer ID), were used to generate cDNAs corresponding to different HIV-1 proteins (RT, PR, IN, Vpr and Env; Supplemental Table 2). Partial HIV-1 genes were then amplified by PCR and sequenced with 300 paired-end Illumina Miseq. A template consensus sequence (TCS 2.0 pipeline, https://primer-id.org/) was generated for sequences with the same Primer ID (i.e. derived from the same viral RNA genome).

Near Full Length (NFL) genome sequencing was performed using previously described methods [5, 10, 11]. Briefly, NFL amplicons were generated by limiting dilution PCR of cDNA or proviral DNA. Second round PCR amplified either a nested NFL amplicon or overlapping 5’ and 3’ half genomes amplicons and sequenced (Supplemental Table 3).

Sequences were aligned using Muscle 5.1 and neighbor-joining phylogenetic trees were generated with Ninja 1.2.2. R5 or X4 tropism were predicted using Geno2Pheno (https://coreceptor.geno2pheno.org/), with a 2% FPR cut-off determining X4-tropic sequences.

### Env Cloning and Entry Phenotype Assays

Full-length *env* amplicons were selected to represent viral diversity within each participant, amplicons were cloned into expression vectors, used to generate luciferase reporter viruses and our previously described method [12] was used to examine the ability of Envs to facilitate efficient entry into Affinofile cells expressing a low surface density of CD4. This assay has previously been shown to distinguish HIV-1 adapted to replication in macrophage (macrophage- or M-tropic) from virus adapted to replication in CD4+ T cells (T-tropic) [13, 14].

### Evolution Analysis

To assess whether ART was able to effectively suppress viral replication, we separately examined whether slow decay lineages were evolving in three participants during ART and before ART. First, for each participant we generated a consensus of partial HIV-1 *env* sequences (V1V3; Primer ID Miseq) in the slow decay lineage present at the earliest on-ART timepoint (ART initiation, P1 and P2; first post-ART timepoint, P4). Second, we separately calculated the pairwise distance (MEGA11) between the consensus sequence and sequences in the slow decay lineage present at all timepoints with >20 sequences. A 2-sided Mann-Whitney test was preformed to compare the average genetic distance at the earliest on-ART timepoint compared to all other timepoints (Graphpad Prism 10) [15]. For one participant (P1) we also performed the analogous assay to assess evolution in the fast decay lineage.

## RESULTS

This study focuses on four participants who initiated ART with very high viral loads, severe immunosuppression and associated coinfections (Table 1; Figure 1A-D). Despite reporting adherence, participants were not virologically suppressed after 24 weeks of ART (Table 1). Adherence was supported by use of supervised ART (P1, P3 and P4), use of long-acting injectable ART (P3 and P4) and a lack of viral evolution during ART (see below). In addition to ART, P3 and P4 were also placed on combination therapies for disseminated *Mycobacterium avium* Complex (dMAC) and other opportunistic infections (Supplemental Figure 1). Despite several changes to these regimens, persistent dMAC bacteremia could not be controlled for P4.

**Figure 1:**
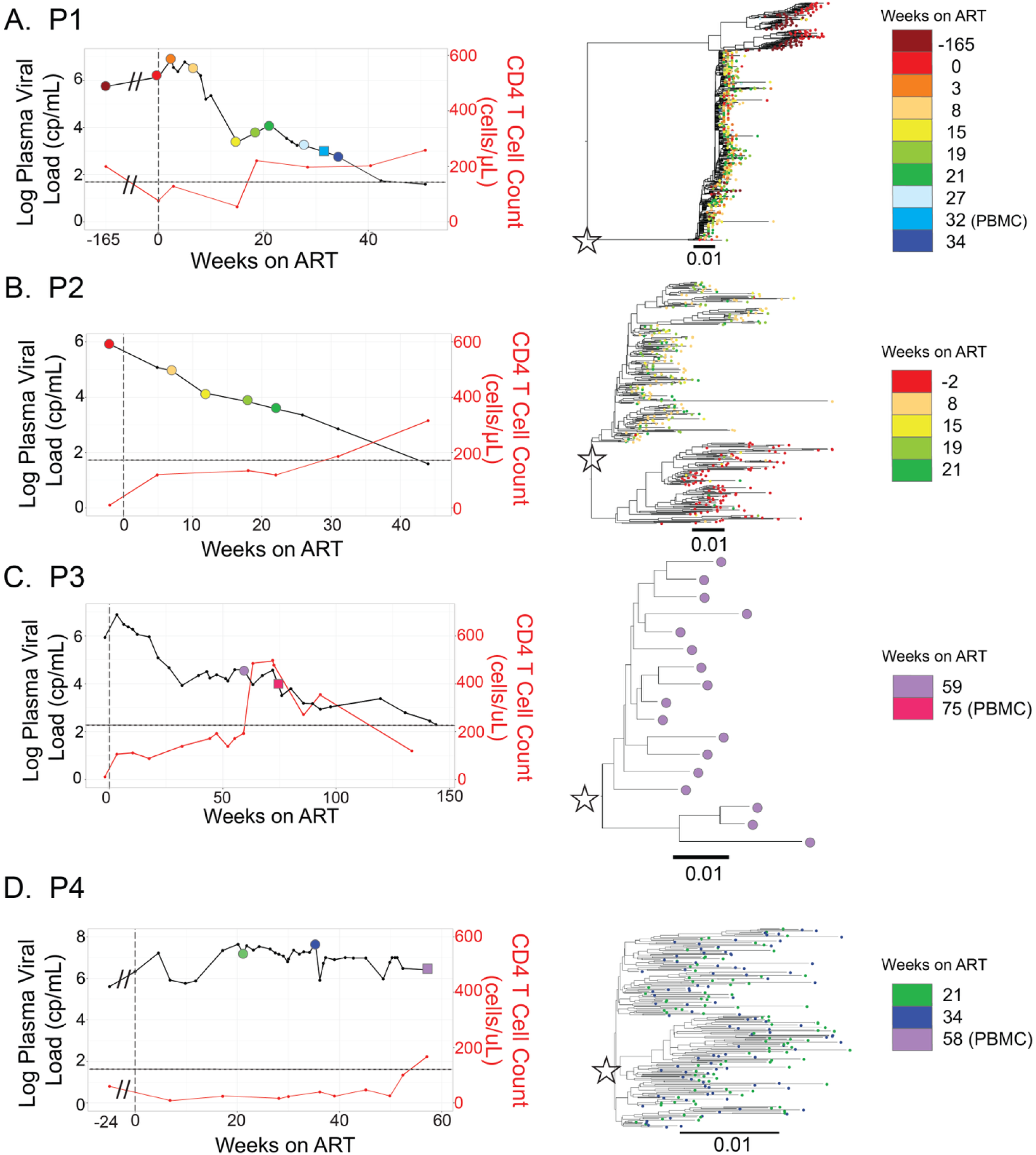
Participants that slowly suppressed viral RNA after initiating ART with drug sensitive virus. Viral loads (HIV-1 RNA cp/mL) and CD4+ T cell (cells/μL; red lines) are shown for all four participants (A-D). Limits of quantification for viral load tests are marked as black, dashed lines. Timepoints when plasma was sampled and used for viral RNA sequencing are represented by circles (⬤) and PBMC samples are represented by squares (▇). Colors indicate when each sample was collected. Phylogenetic trees of partial HIV-1 *env* (V1V3) sequences (identical sequences collapsed) are shown and slow decay lineages are marked with stars.

### Suppression Kinetics and Drug Resistance

After initiating ART, three participants suppressed HIV-1 RNA to undetectable levels in plasma after 50 (P1), 40 (P2) and 144 (P3) weeks (Table 1; Figure 1). For P4, HIV-1 viral load did not decline during the 57 weeks of ART examined by this study, after which the participant died. Deep sequencing was used to assess drug resistance mutations (DRMs) in all participants. No participant had DRMs to their drug regimen that represented more than 1% of the population in their plasma during ART (data not shown).

### HIV-1 Genetic Diversity Prior to ART

Prior to initiating ART, P1 and P2 exhibited highly diverse HIV-1 populations in their plasma. Each had two distinct viral lineages in their blood: one that declined rapidly and another that declined more slowly following ART initiation (Figures 1A-B; Table 2). This pattern is particularly evident in P1, who had both rapid- and slow-decay lineages present at similar frequencies at - 165 and 0 weeks on ART (Table 2). After 8 weeks on ART, the rapid-decay lineage was nearly undetectable, highlighting differential decay kinetics within the viral population. Samples collected proximal to ART initiation were not available for P3 or P4, so we were unable to assess HIV-1 diversity in the plasma before ART.

**Table 2:**
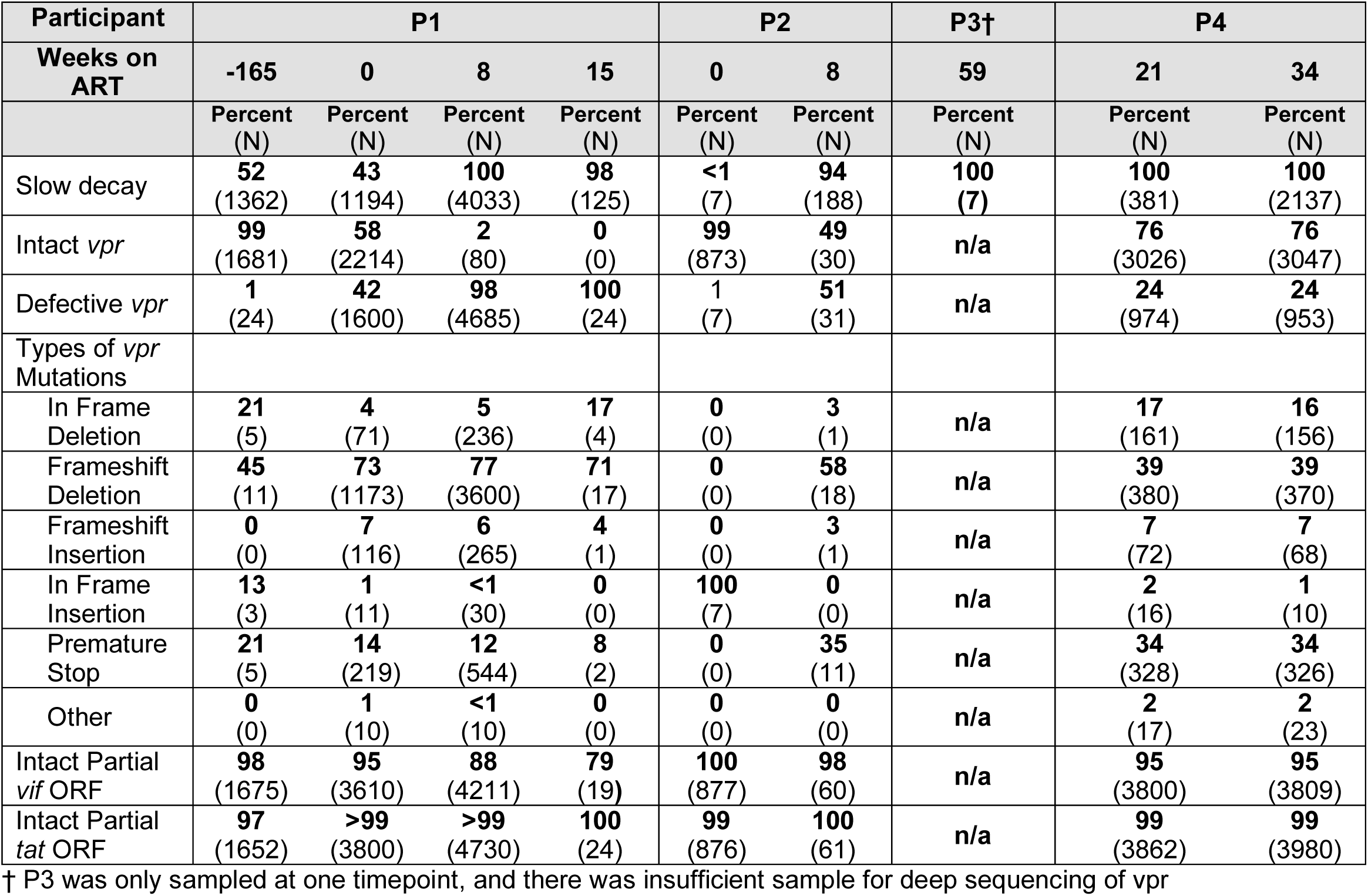
Longitudinal deep sequencing analyses of HIV-1 variants in blood plasma.

### HIV-1 Evolution During ART

Evolution of HIV-1 RNA is an indicator of virus replication. Prior to ART, P1 had two ‘rapid’ decay lineages in their blood (Supplemental Figure 2) as well as a ‘slow’ decay lineage (Figure 1). As expected, analyses revealed that prior to ART both ‘slow’ and ‘rapid’ decay lineages evolved significantly (-165 vs 0 weeks, Supplemental Figure 3A-B). In contrast, analyses of ‘slow’ decay lineages during ART did not find evidence of viral evolution (Supplemental Figure 3B-D). Only one comparison was significant (P1, 0 vs 8 weeks; Supplemental Figure 3B), however, this was driven by the presence of many deletions at one timepoint (week 8). Evolution analyses were not performed for P3 because longitudinal plasma samples were not available.

### Cellular Tropism of Viral RNA and proviral DNA

Viral RNA-derived, full-length *env* amplicons were sequenced for all participants and DNA-derived *envs* were also sequenced for P1, P3 and P4 (Figure 2). All *env* sequences from RNA present in the plasma during ART were predicted to be R5 tropic by Geno2pheno [16] (using an FPR cut off of 2% or less to indicate CXCR4 (X4) -using virus), but some PBMC-derived proviral *env* genes from P4 were predicted to be X4-tropic. Overall, proviral genomes were genetically distinct from viral RNA in the plasma during ART, suggesting that the ‘slow’ decay virus was being produced by cells in another tissue or by a low frequency provirus in the blood. However, P1 had one PBMC-derived provirus that clustered with the ‘slow’ decay RNA lineage.

**Figure 2:**
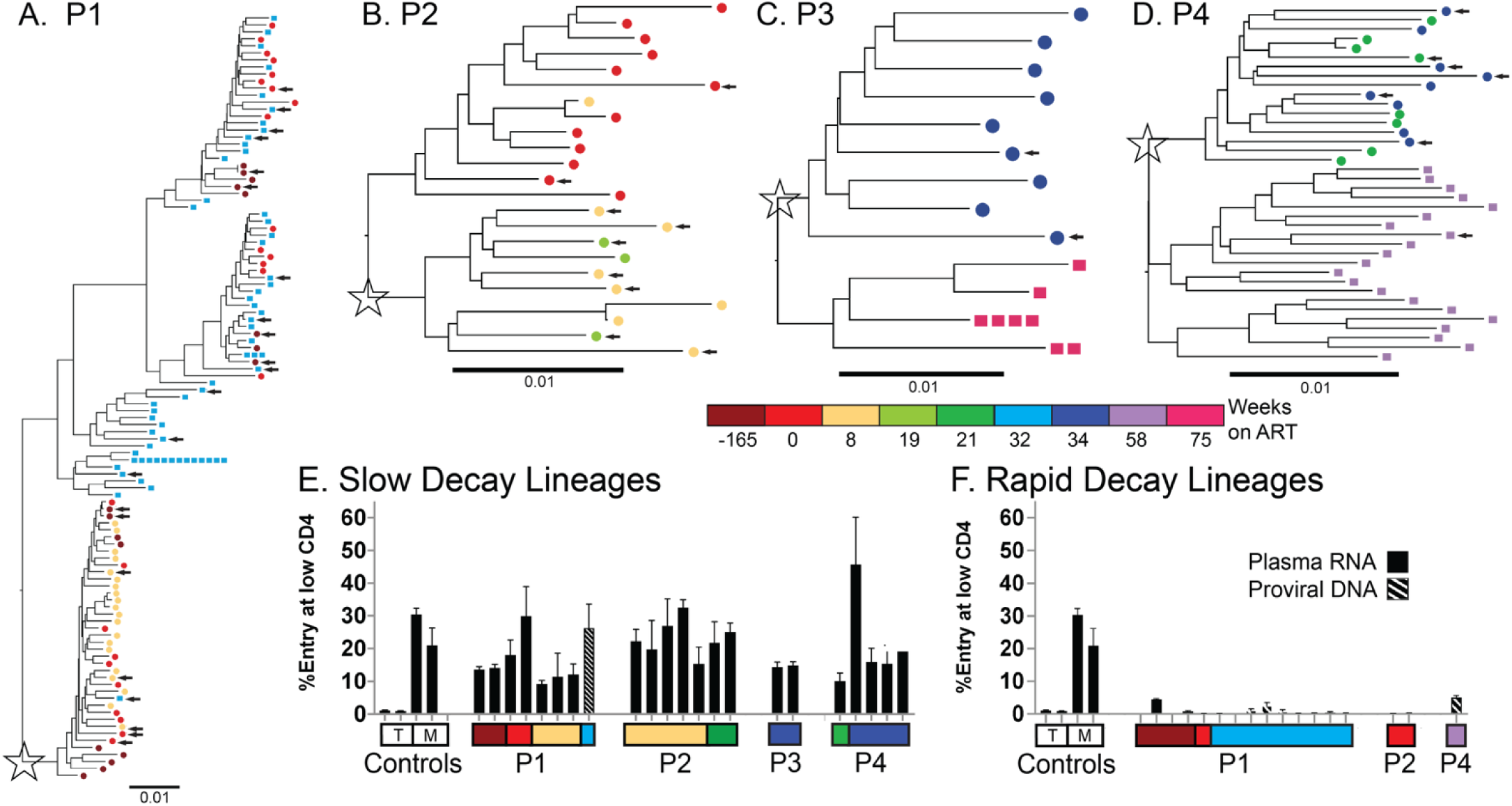
Diversity of intact HIV-1 *env* genes and phenotypic assay for macrophage tropism (M tropism). Phylogenetic trees of intact HIV-1 *env* genes from all Participants are shown (A-D). Amplicons of *envs* were generated at limiting dilution from blood plasma RNA (circles ⬤) and proviral DNA in PBMCs (squares ▇). Plasma lineages that decayed slowly after ART initiation or did not decay are noted with a star and sequences are colored according to their sample time. Luciferase reporter viruses were generated from *envs* (selected *envs* marked with arrows in phylogenetic trees) in both lineages that slowly decayed after ART initiation (E) and lineages that decayed rapidly after ART initiation (F). M tropism was assessed based on the ability of reporter viruses to enter Affinofile cells expressing a low surface density of CD4 (similar to levels on macrophage) relative to their ability to enter Affinofile cells expressing a high density of CD4 (similar to levels on CD4+ T cells). M- and T-tropic controls are shown. The *envs* derived from plasma RNA are shown with solid bars and *envs* from proviral DNA are shown with crosshatched bars.

Intact, full-length, *env* genes representing within participant genetic diversity (Figure 2A-D) were selected and used in entry assays to explore the cellular source of HIV-1 RNA in the plasma during ART. Specifically, whether variants were adapted to replication in CD4+ T cells (T-tropic) or adapted to replication in macrophage (M-tropic). Like our M-tropic controls, *env* genes derived from ‘slow’ decay RNA populations (and one provirus) were all M-tropic based on their ability to efficiently infect cells expressing a low surface density of CD4 (Figure 2E).

In contrast, viral RNA that decayed rapidly after P1 and P2 initiated ART, encoded Env proteins that were unable to efficiently enter cells with a low density of CD4 (Figure 2F), indicating that they were T-tropic. Similarly, analyses of *env* genes from PBMC DNA in P1 found that all but one provirus was both closely related to lineages that decayed rapidly after ART initiation (Figures 2A-B) and T-tropic (Figure 2F). This is consistent with HIV-infected CD4+ T cells being the source of proviral DNA in PBMCs during ART and the source of viral RNA that decays rapidly after initiation of ART. Analyses of a PBMC DNA-derived *env* from P4 also found that it was T-tropic.

### Convergent Evolution of *vpr* Defects in M-tropic Virus

To assess whether additional mutations were linked to virus expression during ART, we performed longitudinal analyses of near full-length genomes from P1 and found that M-tropic (e.g. ‘slow’ decay) viral lineages commonly accumulated defects in *vpr* (Figure 3). No *vpr* mutations were observed in either T- or M-tropic lineages in the plasma 165 weeks prior to ART, or in the T-tropic population at ART initiation. However, 93% (41 of 44) of M-tropic NFL sequences from 0 and 8 weeks on ART had defects in the accessory gene *vpr* (Figure 3).

**Figure 3:**
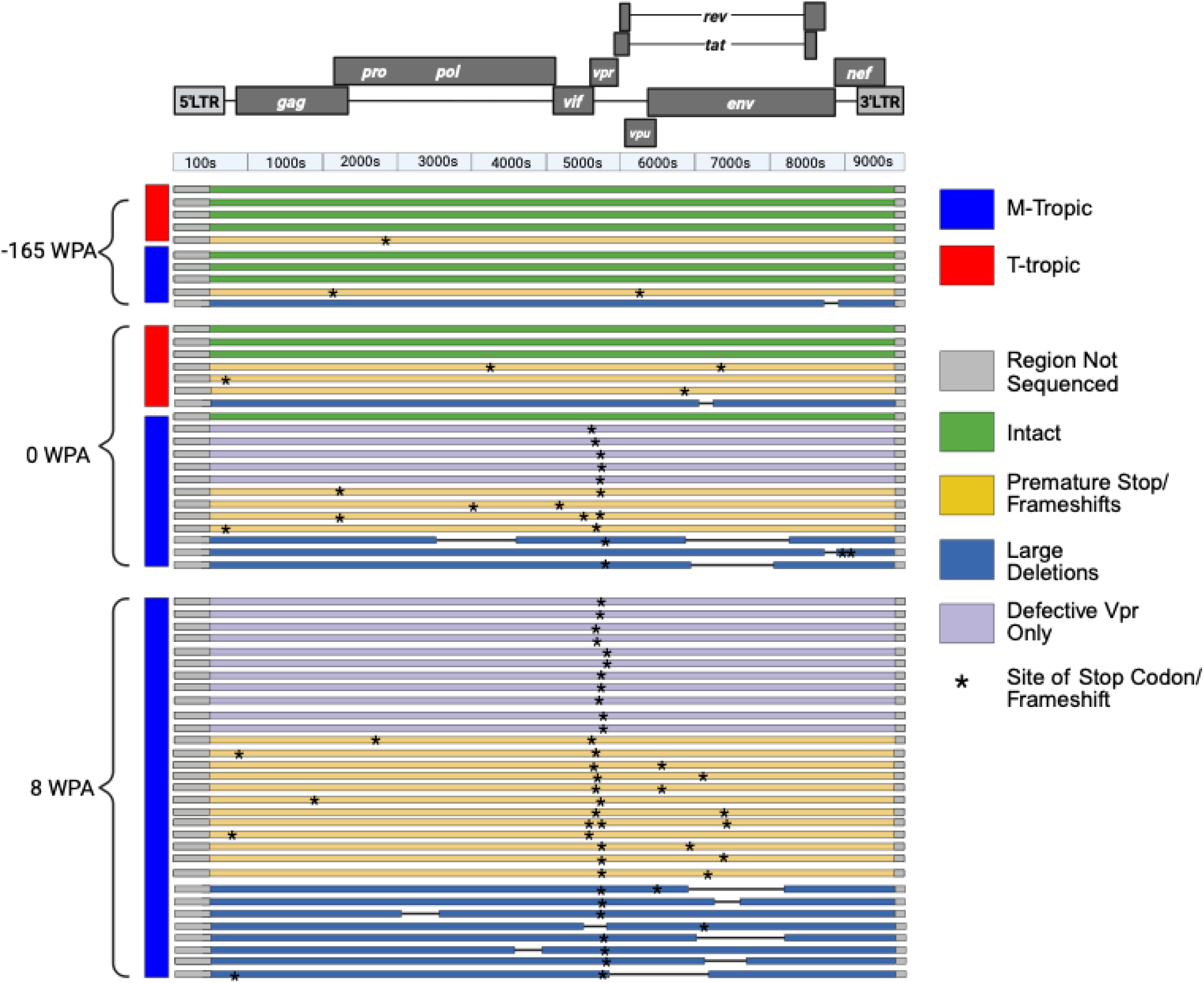
Intactness of near full-length sequences from P1 plasma RNA. Sequences were generated at limiting dilution as two half genomes, and analyzed for intactness of open reading frames, hypermutation, and large deletions. Stars indicate nonsynonymous changes including frameshifts. Sequences are grouped by both their cellular tropism (M-tropic, blue bar on left; T-tropic, red bar on left) and the timepoint when they were sampled. Created in BioRender. Moeser, M. (2025) https://BioRender.com/62gzait.

To better understand the association between defects in *vpr* and M-tropism, we used multiplexed Primer-ID Miseq to generate unlinked *vpr* and partial *env* sequences from P1, P2 and P4. Alignments to full *envs* were used to assess whether partial *envs* were M- or T-tropic and *vpr* sequences were examined for intactness. For the participants with pre- and post-ART plasma samples (P1 and P2), we observed that the proportion of M-tropic/’slow decay’ lineages increased after ART initiation (i.e. after loss of ‘rapid’ decay lineages) and the proportion of *vpr* defects increased in parallel (Table 2 and Figure 4). Analyses of P4 were complicated by the lack of pre-ART sequences and the fact that there was a 2 amino acid deletion in all RNA- and DNA-derived *vpr* sequences (amino acids 86-87). Regardless, 100% of variants in the plasma collected from P4 during ART were M-tropic and at least 25% of *vpr* sequences were defective (Table 2 and Figure 4).

**Figure 4:**
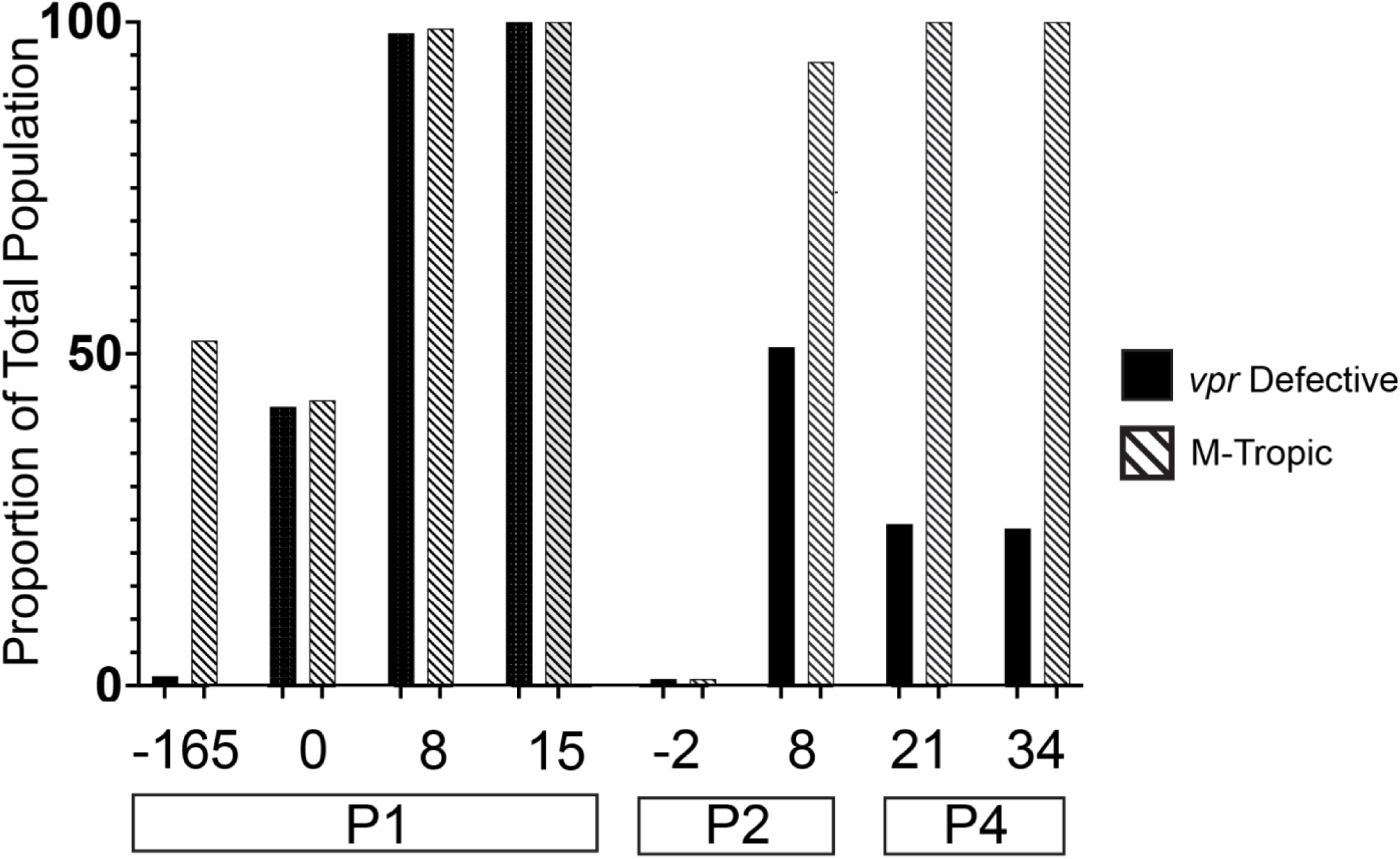
Changes in the frequency of HIV-1 variants with defects in *vpr* and M-tropic HIV-1. Longitudinal deep sequencing of viral RNA in plasma was performed to separately assess *vpr* intactness (solid black bars) and M tropism (crosshatched bars) in participants with longitudinal plasma samples. Proportions determined by Miseq Primer ID sequencing.

We observed many different *vpr* defects, including premature stop codons, frameshifts, and deletions (Table 2 and Supplemental Figure 4). In addition, the *vpr* open reading frame (ORF) partially overlaps with both the *vif* and *tat* ORFs, and we observed that most *vpr* defects accumulated in the nonoverlapping region and 97% of all partial *tat* sequences and 79% of partial *vif* sequences remained intact. These patterns are consistent with *vpr* defects accumulating in replicating virus under conditions where the loss of Vpr protein confers a selective advantage.

## DISCUSSION

In this study, we examined the mechanisms driving persistent HIV-1 viremia in four individuals who initiated ART with advanced disease and high viral loads. While most people with HIV-1 achieve virologic suppression within weeks to months of starting ART, three of our participants required nearly a year to reach suppression and the remaining participant did not achieve suppression after more than a year on ART. Slow (or no) suppression after initiating ART occurred despite some individuals undergoing extensive regimen optimization/intensification and all individuals having both evidence of good ART adherence and drug sensitive virus. Based on these observations we classified these four individuals as having *primary NSV* (i.e. NSV emerging at ART initiation). Additional analyses revealed that these individuals had virus in their plasma during ART that was genetically diverse, not evolving and adapted to replication in macrophage (M-tropic). This indicates that *primary NSV* can be produced by continued expression from HIV-infected macrophage during ART. In contrast, *secondary NSV* (NSV that emerges after a period of viral suppression) has previously been shown to be generated by virus produced from CD4+ T cell clones in people on ART [4–7].

Because HIV-infected macrophage likely reside in tissues and this study was restricted to analyses of blood (plasma and PBMCs), we were unable directly observe HIV-infected macrophage generating *primary NSV*. However, multiple lines of evidence suggest that HIV-infected macrophage were the source of viral RNA during *primary NSV*. First, the viral lineages that persisted during ART were adapted to replication in macrophage. M-tropism is an extremely unusual phenotype that has primarily been observed in the CNS of untreated people with HIV-associated dementia [17] and is very rarely observed in the blood [18]. In contrast, T-tropic HIV-1 (either X4- or R5-using) is the wildtype form of HIV-1 and the type found in the blood of most people [18]. The repeated observation of M-tropic virus in the blood of participants in this study strongly suggests that these populations were being produced by macrophage rather than the typical target cell of HIV-1 (i.e. CD4+ T cells). Further, pre- and post-ART samples were available for two participants and analyses of HIV-1 populations in these samples revealed that M- and T-tropic populations were in the blood before ART, but only the M-tropic lineages persisted during ART. Second, after initiation of ART, viral loads declined with kinetics indicating that they were produced by long-lived cells like macrophage. This differs from the typical pattern in which ART initiation causes viral RNA to rapidly decline, consistent with it being produced by short-lived, CD4+ T cells [19]. Third, the participant who did not suppress HIV-1 RNA during the study period had disseminated Mycobacterium avium complex (MAC) which is thought to enhance HIV-1 infection of macrophage. MAC infects macrophage and *in vitro* analyses have shown that MAC increases both macrophage susceptibility to HIV-1 infection and HIV-1 expression from infected macrophage [20]. Further, *in vivo* mycobacterium infections have been associated with higher HIV-1 viral loads [21] and increased risk for HIV-1 treatment failure [22, 23]. Thus, providing an additional link between *primary NSV* in this person and HIV-1 infection of macrophage. Together, these unique features of participants and viral populations examined in this study are consistent with *primary NSV* being produced by HIV-infected macrophage that continue express virus during ART.

We observed that in three participants, M-tropic lineages independently accumulated many defects in *vpr* while *vpr* genes remained intact in T-tropic lineages observed pre-ART. The accumulation of defects in *vpr* in M-tropic lineages runs counter to the existing literature which suggests that loss of *vpr* inhibits HIV-1 infection of macrophage, but does not impact infection of CD4+ T cells [24–26]. Vpr is a 96 amino acid viral accessory protein that is highly conserved in HIV-1 and HIV-2 [27] and transmission of *vpr* null variants has only been observed four times [28–30]. Vpr is thought to impact many features in host cells including arresting the cell cycle in the G2/M phase [26, 31], increasing genomic instability [32], generating cytotoxicity [33] and increasing transcription from the viral LTR [31]. While *in vitro* studies have reported that *vpr* is necessary for infection of macrophage, but not CD4+ T cells [24–26], our observation of defects in *vpr* in M-tropic lineages in the blood before ART (i.e. in replicating virus) and during ART (i.e. virus being produced from infected macrophage without replication) provides strong evidence that loss of Vpr function impacts both virus replication and expression in macrophage. Possible mechanisms for this advantage include that loss of Vpr function reduces cytotoxicity of HIV-1 in macrophage, inhibits the immune system’s ability to clear HIV-infected macrophage and/or increases HIV-1 expression in infected macrophage. Additional work is needed to better understand the effects of Vpr in macrophage and whether these effects differ before and during ART.

While these people with *primary NSV* initiated ART with severe immunosuppression and high viral loads, it is unclear whether continued HIV-1 production from infected macrophage directly impacted their health. While *secondary NSV* does not have any obvious clinical significance [7], persistent high levels of virus in people with *primary NSV* may continue to drive inflammation and immune dysregulation that could be particularly impactful in the CNS. However, *primary NSV* clearly presented significant challenges for both the participants and the clinicians caring for them, often leading to unnecessary changes in ART or unwarranted diagnostic testing. In addition, we identified some intact viral genomes in the blood during *primary NSV*, raising the possibility that people with *primary NSV* may be able to transmit HIV-1 despite being on ART; a risk that would be amplified by high viral loads in these individuals. Finally, people with *primary NSV* may have large numbers of HIV-infected macrophage that could become latent and contribute to the persistent reservoir. The presence of macrophage reservoirs in people on ART poses specific concerns due to the very long-half of some tissue macrophage [34] and the resistance of HIV-infected macrophage to killing by NK cells [35] and CD8+ T cells [36].

## Acknowledgments

We would like to acknowledge all participants as well as the UNC CFAR Clinical Cohort. We would also like to thank Ronald Swanstrom for helpful discussions.

## Data sharing

All data will be made available at the time of publication.

---

**Supplemental Table 1:**
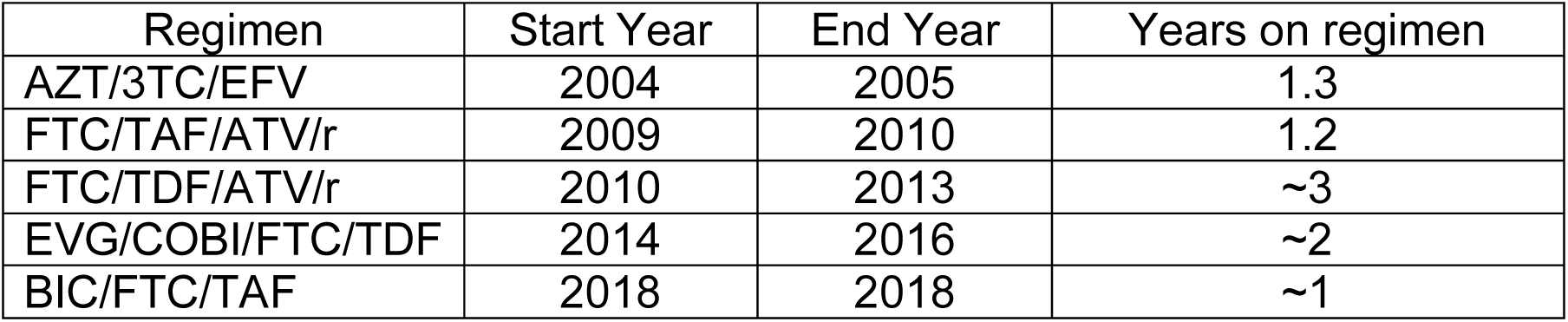
P1 Prior treatment history.

**Supplemental Table 2:**
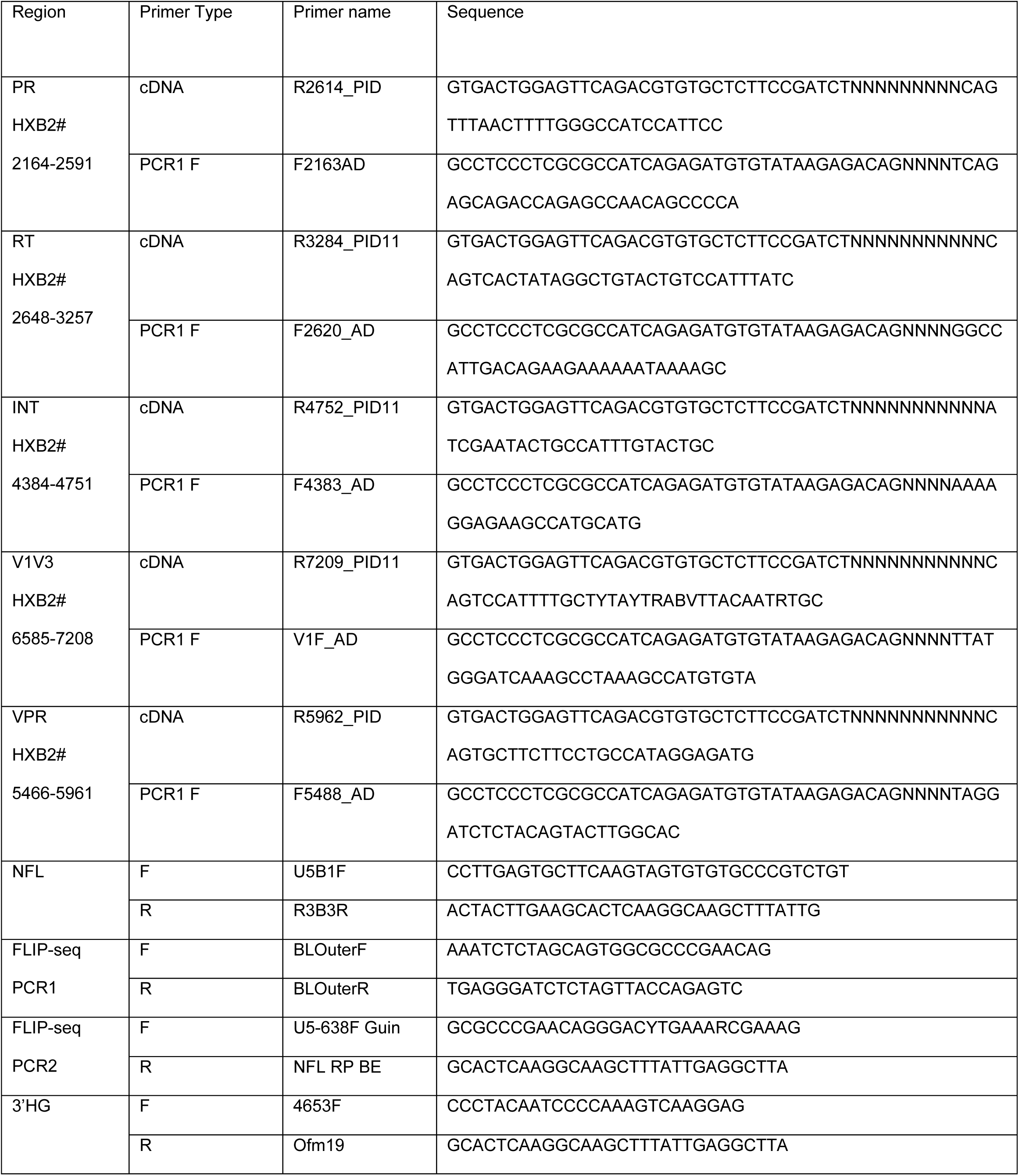
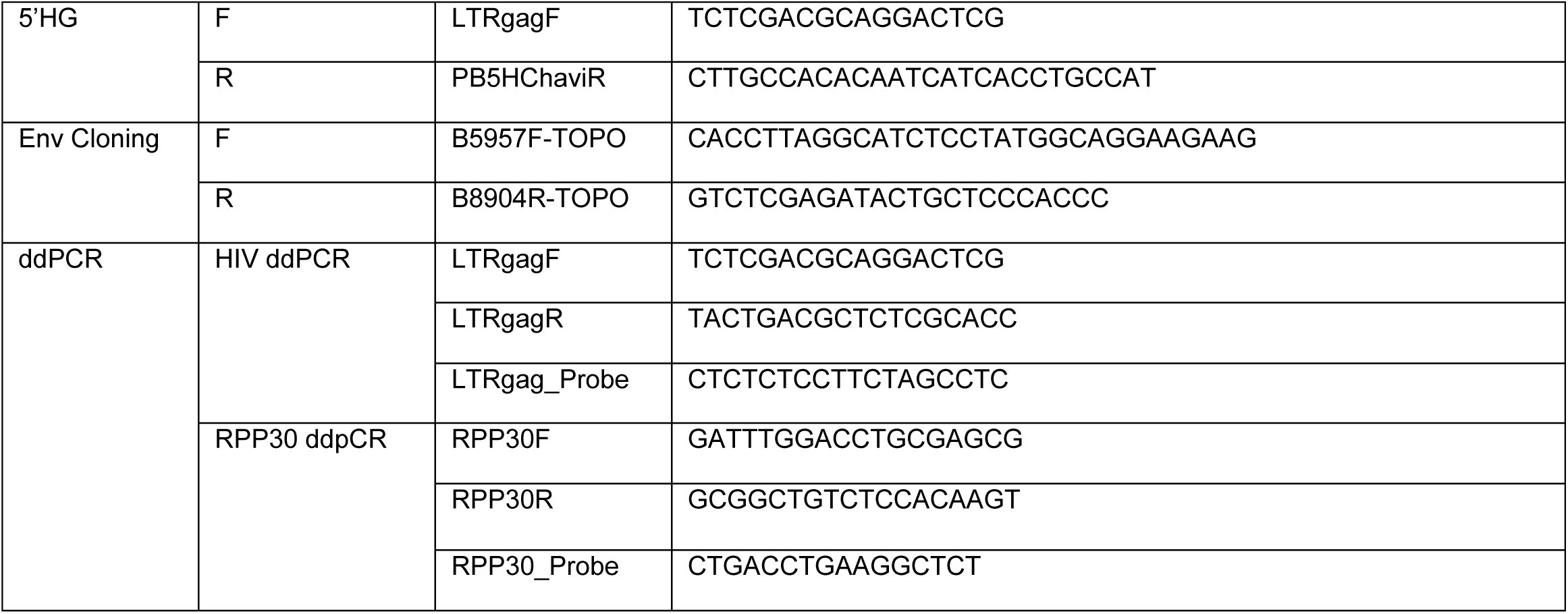
Primers.

**Supplemental Table 3:**
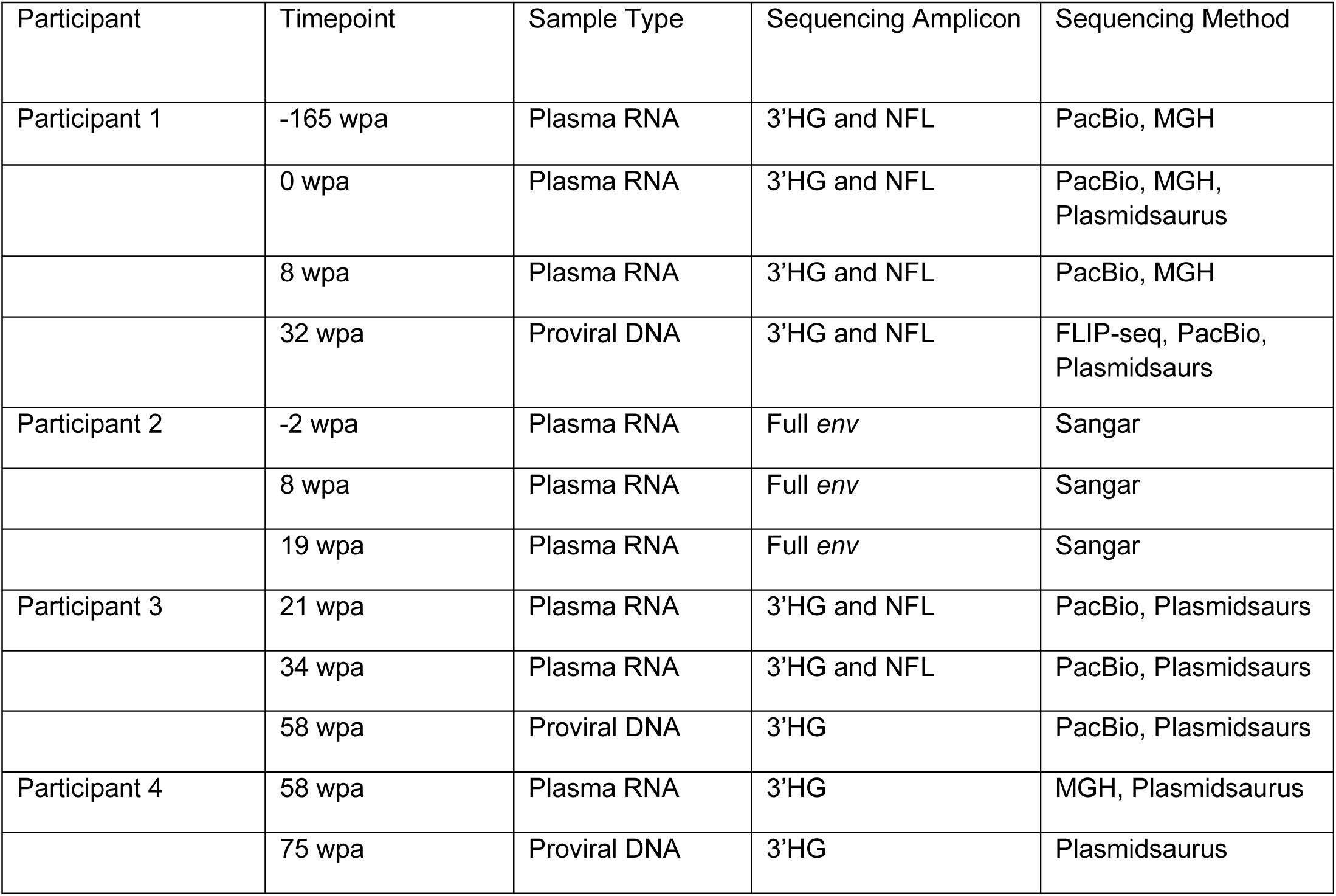
Limiting dilution sequencing methods.

**Supplemental Figure 1:**
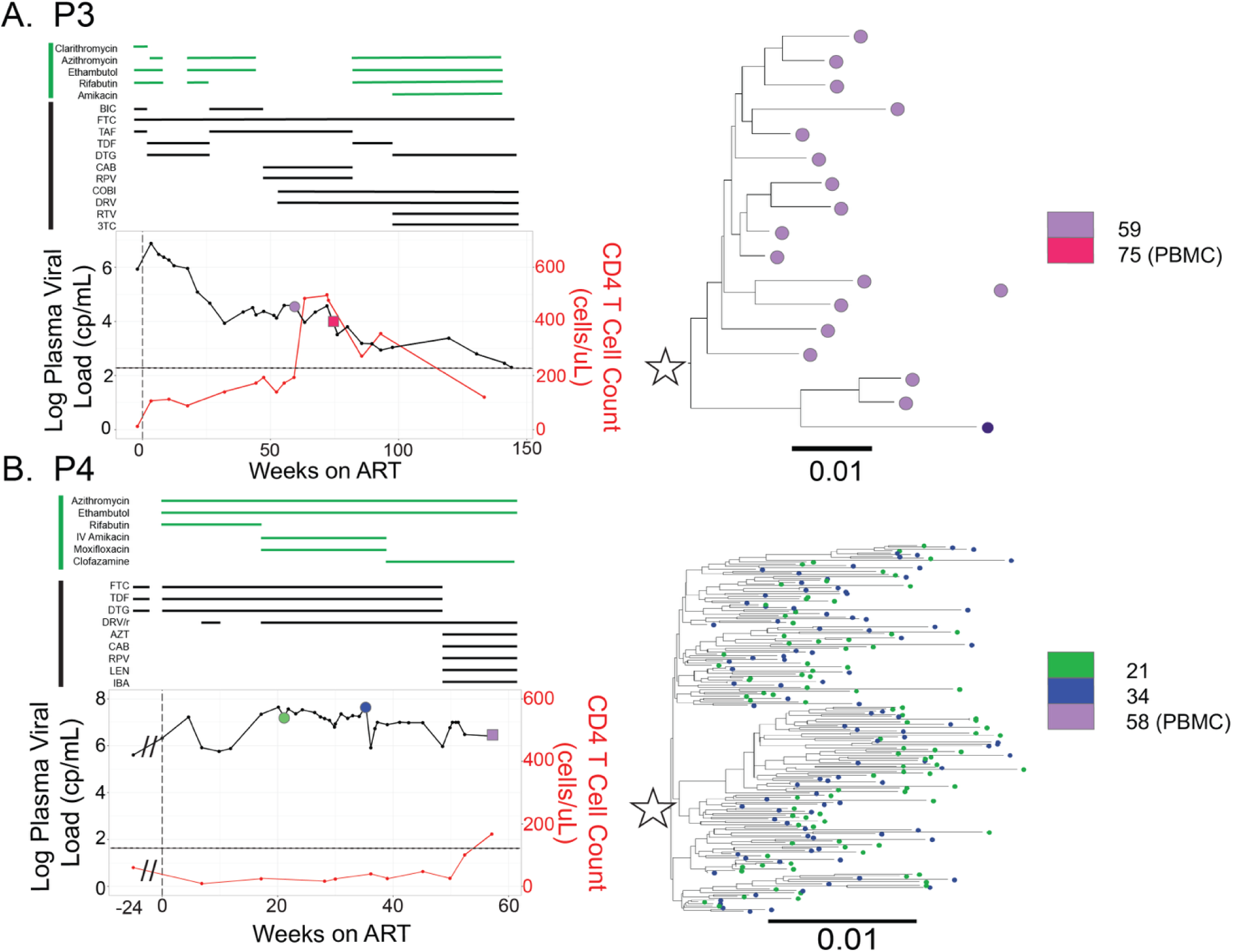
Detailed treatment histories for P3 and P4. (A-B) Viral loads (HIV-1 RNA cp/mL) and CD4+ T cell in cells/μL for P3 and P4. Limits of quantification for viral load tests are marked as a black, dashed lines. Timepoints when plasma was sampled and used for viral RNA sequencing are represented by circles (⬤) and PBMC samples are represented by squares (▇). Colors indicate when each sample was collected. Phylogenetic trees of partial HIV-1 *env* (V1V3) sequences (identical sequences collapsed) are shown and slow decay lineages are marked with stars.

**Supplement Figure 2:**
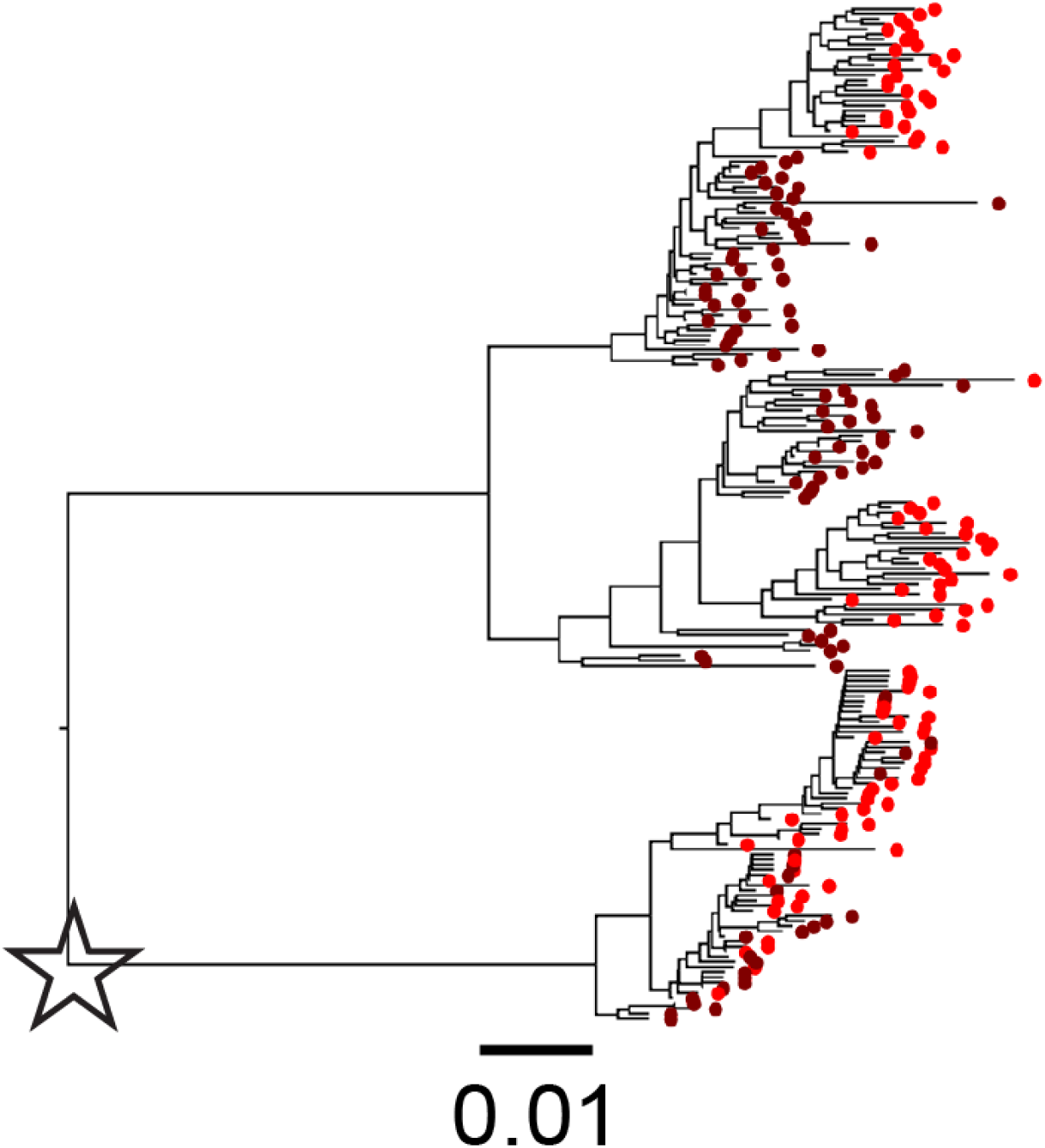
Phylogenetic tree analyzing pre-ART HIV-1 RNA sequences from P1. Phylogenetic tree of partial HIV-1 *env* (V1V3) sequences (identical sequences collapsed) of viral RNA in plasma collected at 2 timepoints before ART initiation (see timepoints and colors in 1A) and the lineage that persists after ART is noted with a star.

**Supplemental Figure 3:**
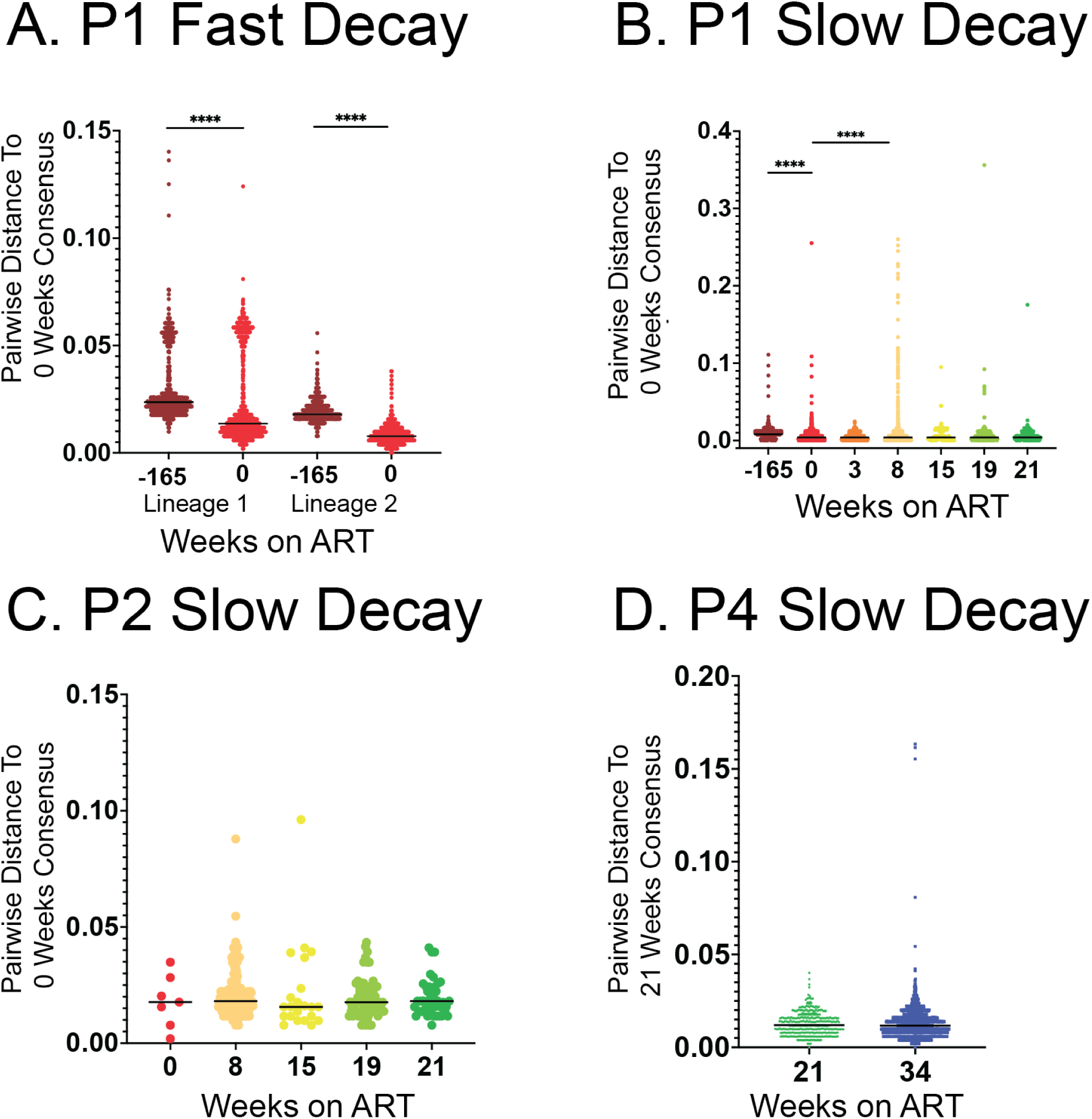
Evolution during ART. A consensus sequence was generated from partial *env* sequences produced from viral RNA in the Pre-ART timepoint population for P1 (A, B) and P2 (C), and the earliest available viral RNA timepoint for P4. Pairwise distance from the consensus was calculated to every sequence at all plasma timepoints with >20 sequences. The pairwise distance populations of the Pre-ART timepoint (P1, P2) and earliest timepoint (P4) were compared to all other timepoints with a Mann-Witney test with a Bonferroni correction.

**Supplement Figure 4:**
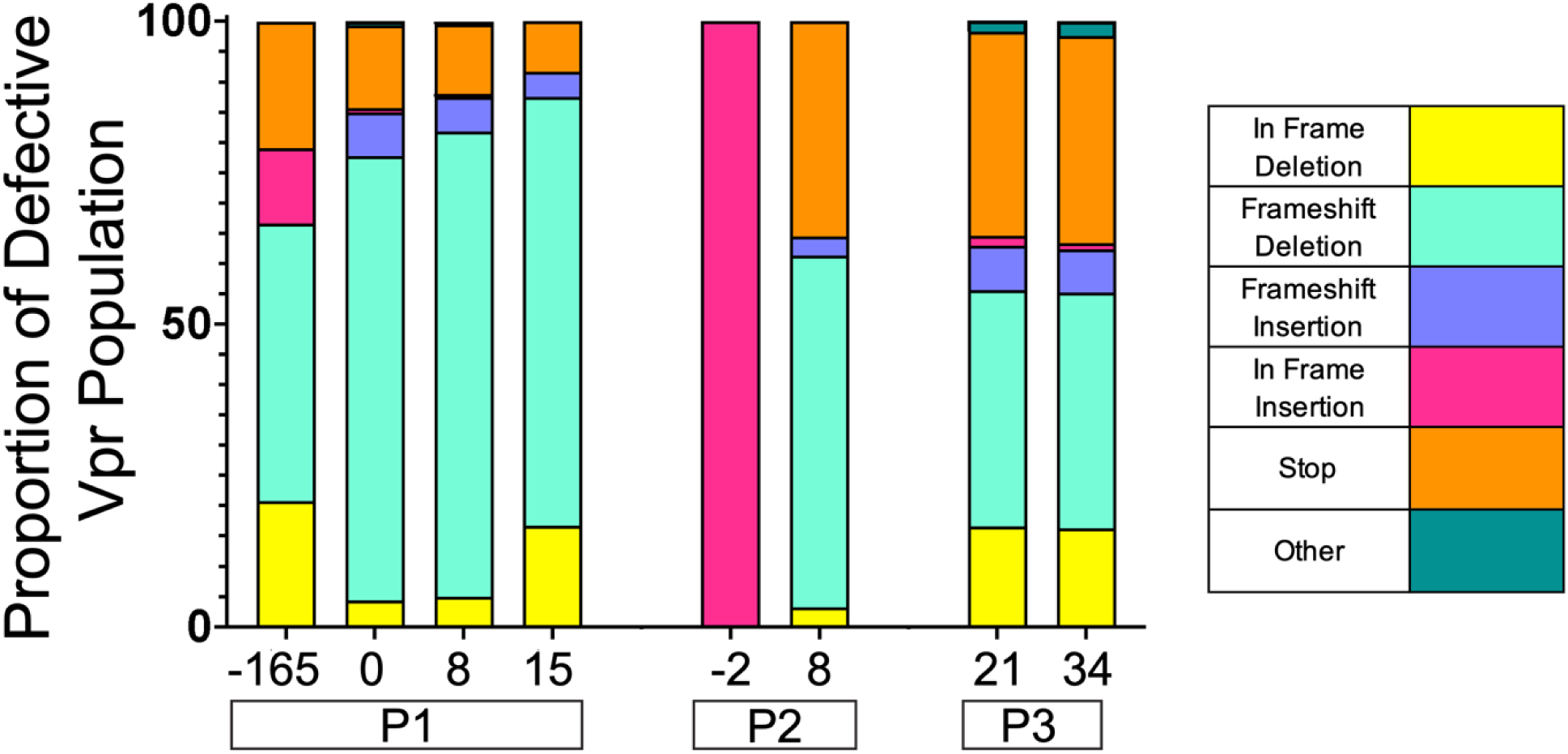
Proportion of *vpr* defects within the entire defective *vpr* gene population at each sampled timepoint. Proportions determined by Miseq Primer ID sequencing.

## REFERENCES

1. Gandhi, R.T., et al., Antiretroviral Drugs for Treatment and Prevention of HIV in Adults: 2024 Recommendations of the International Antiviral Society-USA Panel. JAMA, 2025. 333(7): p. 609–628.

2. Hamers, R.L., et al., Effect of pretreatment HIV-1 drug resistance on immunological, virological, and drug-resistance outcomes of first-line antiretroviral treatment in sub-Saharan Africa: a multicentre cohort study. Lancet Infect Dis, 2012. 12(4): p. 307–17.

3. Pinoges, L., et al., Risk factors and mortality associated with resistance to first-line antiretroviral therapy: multicentric cross-sectional and longitudinal analyses. J Acquir Immune Defic Syndr, 2015. 68(5): p. 527–35.

4. Halvas, E.K., et al., HIV-1 viremia not suppressible by antiretroviral therapy can originate from large T cell clones producing infectious virus. J Clin Invest, 2020. 130(11): p. 5847–5857.

5. Mohammadi, A., et al., Viral and host mediators of non-suppressible HIV-1 viremia. Nat Med, 2023.

6. Simonetti, F.R., et al., Clonally expanded CD4+ T cells can produce infectious HIV-1 in vivo. Proc Natl Acad Sci U S A, 2016. 113(7): p. 1883–8.

7. White, J.A., et al., Clonally expanded HIV-1 proviruses with 5’-leader defects can give rise to nonsuppressible residual viremia. J Clin Invest, 2023. 133(6).

8. Joseph, S.B., et al., Human Immunodeficiency Virus Type 1 RNA Detected in the Central Nervous System (CNS) After Years of Suppressive Antiretroviral Therapy Can Originate from a Replicating CNS Reservoir or Clonally Expanded Cells. Clin Infect Dis, 2019. 69(8): p. 1345–1352.

9. Zhou, S., et al., Primer ID Validates Template Sampling Depth and Greatly Reduces the Error Rate of Next-Generation Sequencing of HIV-1 Genomic RNA Populations. J Virol, 2015. 89(16): p. 8540–55.

10. Joseph, S.B., et al., The timing of HIV-1 infection of cells that persist on therapy is not strongly influenced by replication competency or cellular tropism of the provirus. PLoS Pathog, 2024. 20(2): p. e1011974.

11. Abrahams, M.R.J., S. B.; Garrett, N.; Tyers, L; Moeser, M; Archin, N; Council, O. D.; Matten, D; Zhou, S; Doolabh, D; Anthony, C; Goonetilleke, N; Karim, S. A.; Margolis, D. M.; Pond, S. K.; williamson, C; Swanstrom, R., The replication-competent HIV-1 latent reservoir is primarily established near the time of therapy initiation. Science Translational Medicine, 2019. 11.

12. Joseph, S.B., B. Lee, and R. Swanstrom, Affinofile Assay for Identifying Macrophage-Tropic HIV-1. Bio Protoc, 2014. 4(14).

13. Johnston, S.H., et al., A quantitative affinity-profiling system that reveals distinct CD4/CCR5 usage patterns among human immunodeficiency virus type 1 and simian immunodeficiency virus strains. J Virol, 2009. 83(21): p. 11016–26.

14. Joseph, S.B., et al., Quantification of entry phenotypes of macrophage-tropic HIV-1 across a wide range of CD4 densities. J Virol, 2014. 88(4): p. 1858–69.

15. Lee, S.K., et al., Sequence Analysis of Inducible, Replication-Competent Virus Reveals No Evidence of HIV-1 Evolution During Suppressive Antiviral Therapy, Indicating a Lack of Ongoing Viral Replication. Open Forum Infect Dis, 2024. 11(5): p. ofae212.

16. Lengauer, T., et al., Bioinformatics prediction of HIV coreceptor usage. Nat Biotechnol, 2007. 25(12): p. 1407–10.

17. Joseph, S.B. and R. Swanstrom, The evolution of HIV-1 entry phenotypes as a guide to changing target cells. J Leukoc Biol, 2018. 103(3): p. 421–431.

18. Ping, L.H., et al., Comparison of viral Env proteins from acute and chronic infections with subtype C human immunodeficiency virus type 1 identifies differences in glycosylation and CCR5 utilization and suggests a new strategy for immunogen design. J Virol, 2013. 87(13): p. 7218–33.

19. Perelson, A.S., et al., HIV-1 dynamics in vivo: virion clearance rate, infected cell life-span, and viral generation time. Science, 1996. 271(5255): p. 1582–6.

20. Wahl, S.M., et al., Mycobacterium avium complex augments macrophage HIV-1 production and increases CCR5 expression. Proc Natl Acad Sci U S A, 1998. 95(21): p. 12574–9.

21. Donovan, R.M., et al., Changes in virus load markers during AIDS-associated opportunistic diseases in human immunodeficiency virus-infected persons. J Infect Dis, 1996. 174(2): p. 401–3.

22. Richterman, A., et al., Predictors of Clinical Outcomes among People with HIV and Tuberculosis Symptoms after Rapid Treatment Initiation in Haiti. medRxiv, 2024.

23. Afrane, A.K.A., et al., HIV virological non-suppression and its associated factors in children on antiretroviral therapy at a major treatment centre in Southern Ghana: a cross-sectional study. BMC Infect Dis, 2021. 21(1): p. 731.

24. Connor, R.I., et al., Vpr is required for efficient replication of human immunodeficiency virus type-1 in mononuclear phagocytes. Virology, 1995. 206(2): p. 935–44.

25. Mashiba, M., et al., Vpr overcomes macrophage-specific restriction of HIV-1 Env expression and virion production. Cell Host Microbe, 2014. 16(6): p. 722–35.

26. Zhang, F. and P.D. Bieniasz, HIV-1 Vpr induces cell cycle arrest and enhances viral gene expression by depleting CCDC137. Elife, 2020. 9.

27. Vanegas-Torres, C.A. and M. Schindler, HIV-1 Vpr Functions in Primary CD4(+) T Cells. Viruses, 2024. 16(3).

28. Ali, A., et al., Highly Attenuated Infection With a Vpr-Deleted Molecular Clone of Human Immunodeficiency Virus-1. J Infect Dis, 2018. 218(9): p. 1447–1452.

29. Beaumont, T., et al., Reversal of human immunodeficiency virus type 1 IIIB to a neutralization-resistant phenotype in an accidentally infected laboratory worker with a progressive clinical course. J Virol, 2001. 75(5): p. 2246–52.

30. Wang, B., et al., Gene defects clustered at the C-terminus of the vpr gene of HIV-1 in long-term nonprogressing mother and child pair: in vivo evolution of vpr quasispecies in blood and plasma. Virology, 1996. 223(1): p. 224–32.

31. Goh, W.C., et al., HIV-1 Vpr increases viral expression by manipulation of the cell cycle: a mechanism for selection of Vpr in vivo. Nat Med, 1998. 4(1): p. 65–71.

32. Laguette, N., et al., Premature activation of the SLX4 complex by Vpr promotes G2/M arrest and escape from innate immune sensing. Cell, 2014. 156(1-2): p. 134–45.

33. Somasundaran, M., et al., Evidence for a cytopathogenicity determinant in HIV-1 Vpr. Proc Natl Acad Sci U S A, 2002. 99(14): p. 9503–8.

34. Reu, P., et al., The Lifespan and Turnover of Microglia in the Human Brain. Cell Rep, 2017. 20(4): p. 779–784.

35. Clayton, K.L., et al., HIV-infected macrophages resist efficient NK cell-mediated killing while preserving inflammatory cytokine responses. Cell Host Microbe, 2021. 29(3): p. 435–447 e9.

36. Clayton, K.L., et al., Resistance of HIV-infected macrophages to CD8(+) T lymphocyte-mediated killing drives activation of the immune system. Nat Immunol, 2018. 19(5): p. 475–486.

