## Supplemental Table 1 for "A New Type of Nonsuppressible Viremia Produced by HIV-Infected Macrophage"

**Supplemental Table 1:** P1 Prior treatment history

| Regimen | Start Year | End Year | Years on regimen |
|---|---|---|---|
| AZT/3TC/EFV | 2004 | 2005 | 1.3 |
| FTC/TAF/ATV/r | 2009 | 2010 | 1.2 |
| FTC/TDF/ATV/r | 2010 | 2013 | ~3 |
| EVG/COBI/FTC/TDF | 2014 | 2016 | ~2 |
| BIC/FTC/TAF | 2018 | 2018 | ~1 |
