## Supplemental Table 2 for "A New Type of Nonsuppressible Viremia Produced by HIV-Infected Macrophage"

**Supplemental Table 2:** Primers

| Region | Primer Type | Primer name | Sequence |
|---|---|---|---|
| PR HXB2# 2164-2591 | cDNA | R2614_PID | GTGACTGGAGTTCAGACGTGTGCTCTTCCGATCTNNNNNNNNNCAG TTTAACTTTTGGGCCATCCATTCC |
| | PCR1 F | F2163AD | GCCTCCCTCGCGCCATCAGAGATGTGTATAAGAGACAGNNNNTCAG AGCAGACCAGAGCCAACAGCCCCA |
| RT HXB2# 2648-3257 | cDNA | R3284_PID11 | GTGACTGGAGTTCAGACGTGTGCTCTTCCGATCTNNNNNNNNNNNC AGTCACTATAGGCTGTACTGTCCATTTATC |
| | PCR1 F | F2620_AD | GCCTCCCTCGCGCCATCAGAGATGTGTATAAGAGACAGNNNNGGCC ATTGACAGAAGAAAAAATAAAAGC |
| INT HXB2# 4384-4751 | cDNA | R4752_PID11 | GTGACTGGAGTTCAGACGTGTGCTCTTCCGATCTNNNNNNNNNNNA TCGAATACTGCCATTTGTACTGC |
| | PCR1 F | F4383_AD | GCCTCCCTCGCGCCATCAGAGATGTGTATAAGAGACAGNNNNAAAA GGAGAAGCCATGCATG |
| V1V3 HXB2# 6585-7208 | cDNA | R7209_PID11 | GTGACTGGAGTTCAGACGTGTGCTCTTCCGATCTNNNNNNNNNNNC AGTCCATTTTGCTYTAYTRABVTTACAATRTGC |
| | PCR1 F | V1F_AD | GCCTCCCTCGCGCCATCAGAGATGTGTATAAGAGACAGNNNNTTAT GGGATCAAAGCCTAAAGCCATGTGTA |
| VPR HXB2# 5466-5961 | cDNA | R5962_PID | GTGACTGGAGTTCAGACGTGTGCTCTTCCGATCTNNNNNNNNNNNC AGTGCTTCTTCCTGCCATAGGAGATG |
| | PCR1 F | F5488_AD | GCCTCCCTCGCGCCATCAGAGATGTGTATAAGAGACAGNNNNTAGG ATCTCTACAGTACTTGGCAC |
| NFL | F | U5B1F | CCTTGAGTGCTTCAAGTAGTGTGTGCCCGTCTGT |
| | R | R3B3R | ACTACTTGAAGCACTCAAGGCAAGCTTTATTG |
| FLIP-seq PCR1 | F | BLOuterF | AAATCTCTAGCAGTGGCGCCCGAACAG |
| | R | BLOuterR | TGAGGGATCTCTAGTTACCAGAGTC |
| FLIP-seq PCR2 | F | U5-638F Guin | GCGCCCGAACAGGGACYTGAAARCGAAAG |
| | R | NFL RP BE | GCACTCAAGGCAAGCTTTATTGAGGCTTA |
| 3'HG | F | 4653F | CCCTACAATCCCCAAAGTCAAGGAG |
| | R | Ofm19 | GCACTCAAGGCAAGCTTTATTGAGGCTTA |
| 5'HG | F | LTRgagF | TCTCGACGCAGGACTCG |
| | R | PB5HChaviR | CTTGCCACACAATCATCACCTGCCAT |
| Env Cloning | F | B5957F-TOPO | CACCTTAGGCATCTCCTATGGCAGGAAGAAG |
