## Supplemental Table 3 for "A New Type of Nonsuppressible Viremia Produced by HIV-Infected Macrophage"

| | | | |
|---|---|---|---|
| | R | B8904R-TOPO | GTCTCGAGATACTGCTCCCACCC |
| ddPCR | HIV ddPCR | LTRgagF | TCTCGACGCAGGACTCG |
| | | LTRgagR | TACTGACGCTCTCGCACC |
| | | LTRgag_Probe | CTCTCTCCTTCTAGCCTC |
| | RPP30 ddpCR | RPP30F | GATTTGGACCTGCGAGCG |
| | | RPP30R | GCGGCTGTCTCCACAAGT |
| | | RPP30_Probe | CTGACCTGAAGGCTCT |
