## Supplemental Table 4 for "A New Type of Nonsuppressible Viremia Produced by HIV-Infected Macrophage"

**Supplemental Table 3:** List of limiting dilution sequencing methods

| Participant | Timepoint | Sample Type | Sequencing Amplicon | Sequencing Method |
|---|---|---|---|---|
| Participant 1 | -165 wpa | Plasma RNA | 3’HG and NFL | PacBio, MGH |
| | 0 wpa | Plasma RNA | 3’HG and NFL | PacBio, MGH, Plasmidsaurus |
| | 8 wpa | Plasma RNA | 3’HG and NFL | PacBio, MGH |
| | 32 wpa | Proviral DNA | 3’HG and NFL | FLIP-seq, PacBio, Plasmidsaurs |
| Participant 2 | -2 wpa | Plasma RNA | Full *env* | Sangar |
| | 8 wpa | Plasma RNA | Full *env* | Sangar |
| | 19 wpa | Plasma RNA | Full *env* | Sangar |
| Participant 3 | 21 wpa | Plasma RNA | 3’HG and NFL | PacBio, Plasmidsaurs |
| | 34 wpa | Plasma RNA | 3’HG and NFL | PacBio, Plasmidsaurs |
| | 58 wpa | Proviral DNA | 3’HG | PacBio, Plasmidsaurs |
| Participant 4 | 58 wpa | Plasma RNA | 3’HG | MGH, Plasmidsaurus |
| | 75 wpa | Proviral DNA | 3’HG | Plasmidsaurus |
