## Supplemental Figure 1 for "A New Type of Nonsuppressible Viremia Produced by HIV-Infected Macrophage"

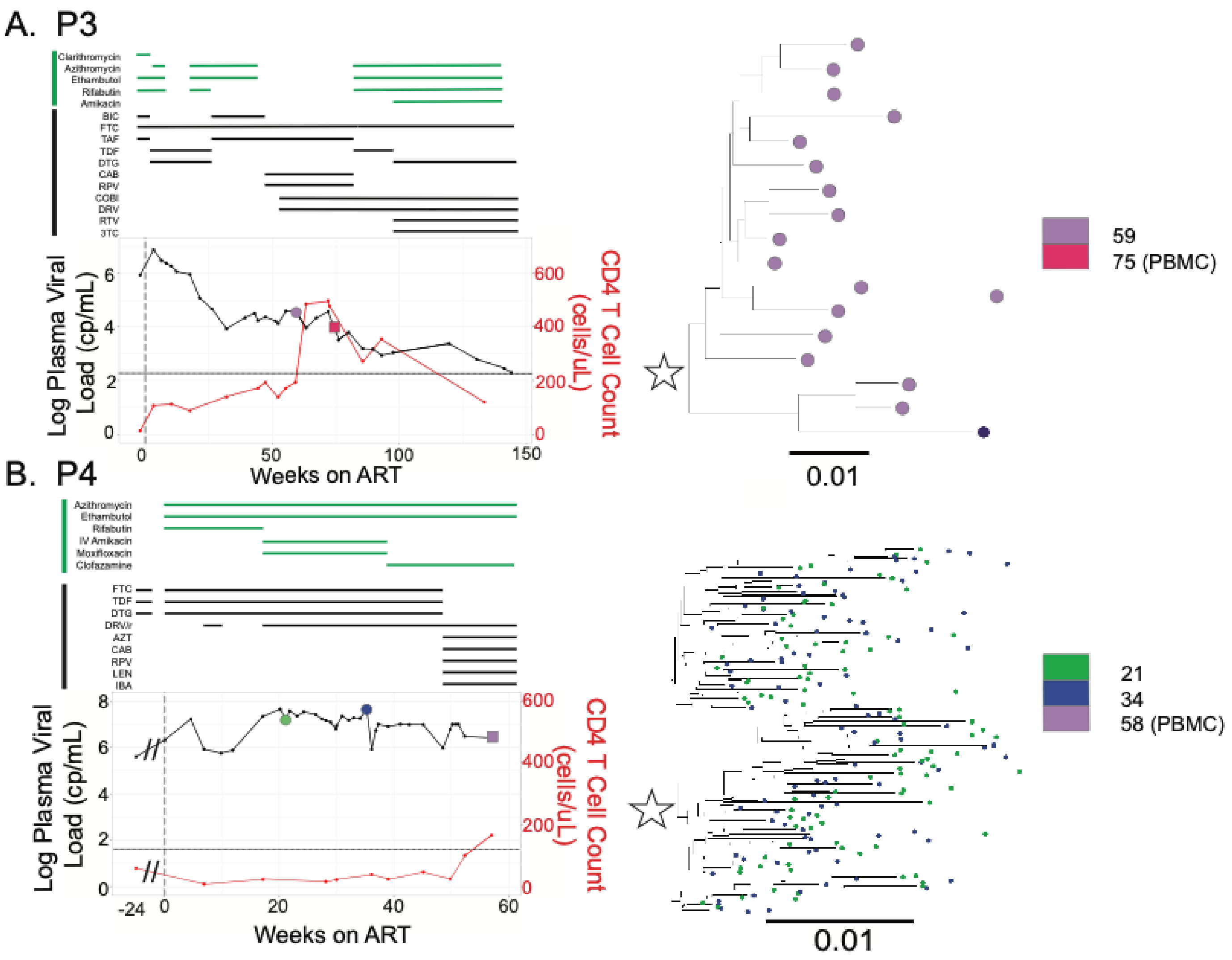


**Supplemental Figure 1: Detailed treatment histories for P3 and P4.** (A-B) Viral loads (HIV-1 RNA cp/mL) and CD4+ T cell in cells/µL for P3 and P4. Limits of quantification for viral load tests are marked as a black, dashed lines. Timepoints when plasma was sampled and used for viral RNA sequencing are represented by circles (●) and PBMC samples are represented by squares (■). Colors indicate when each sample was collected. Phylogenetic trees of partial HIV-1 *env* (V1V3) sequences (identical sequences collapsed) are shown and slow decay lineages are marked with stars.
