## Supplemental Figure 2 for "A New Type of Nonsuppressible Viremia Produced by HIV-Infected Macrophage"

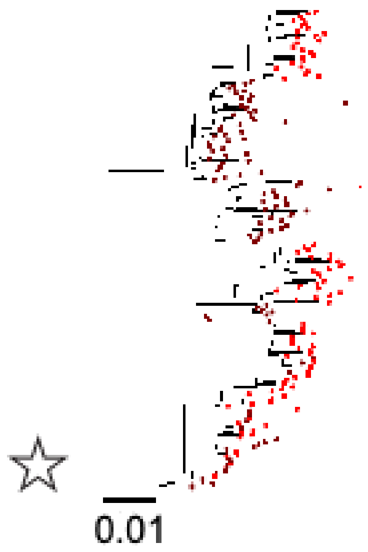


**Supplement Figure 2:** Phylogenetic tree analyzing pre-ART HIV-1 RNA sequences from Participant 1. Phylogenetic tree of partial HIV-1 *env* (V1V3) sequences (identical sequences collapsed) of viral RNA in plasma collected at 2 timepoints before ART initiation (see timepoints and colors in 1A) and the lineage that persists after ART is noted with a star.
