## Supplemental Figure 3 for "A New Type of Nonsuppressible Viremia Produced by HIV-Infected Macrophage"

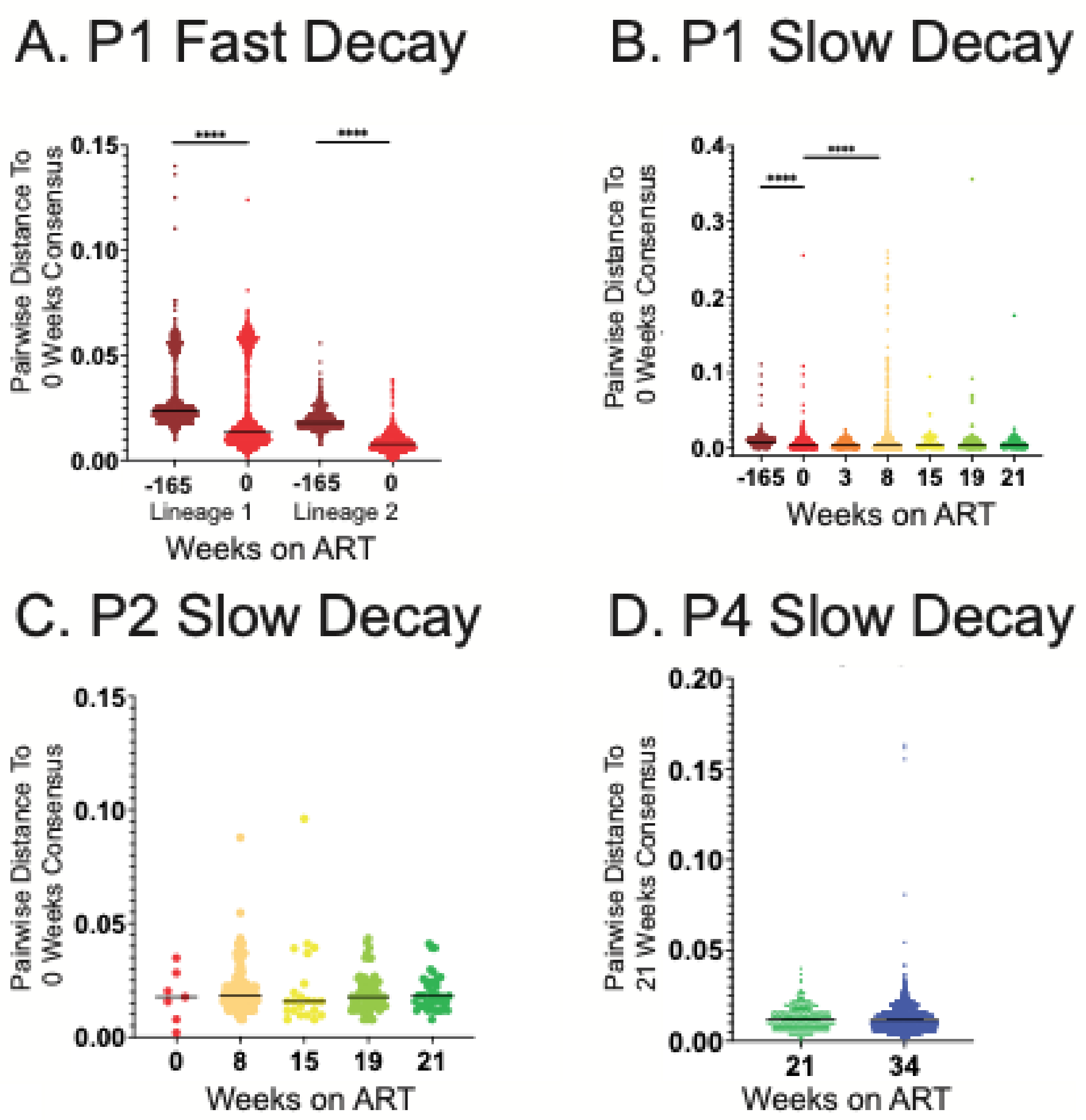


**Supplemental Figure 3: Evolution during ART.** A consensus sequence was generated from partial *env* sequences produced from viral RNA in the Pre-ART timepoint population for P1 (A, B) and P2 (C), and the earliest available viral RNA timepoint for P4. Pairwise distance from the consensus was calculated to every sequence at all plasma timepoints with >20 sequences. The pairwise distance populations of the Pre-ART timepoint (P1, P2) and earliest timepoint (P4) were compared to all other timepoints with a Mann-Witney test with a Bonferrari correction.
