## Supplemental Figure 4 for "A New Type of Nonsuppressible Viremia Produced by HIV-Infected Macrophage"

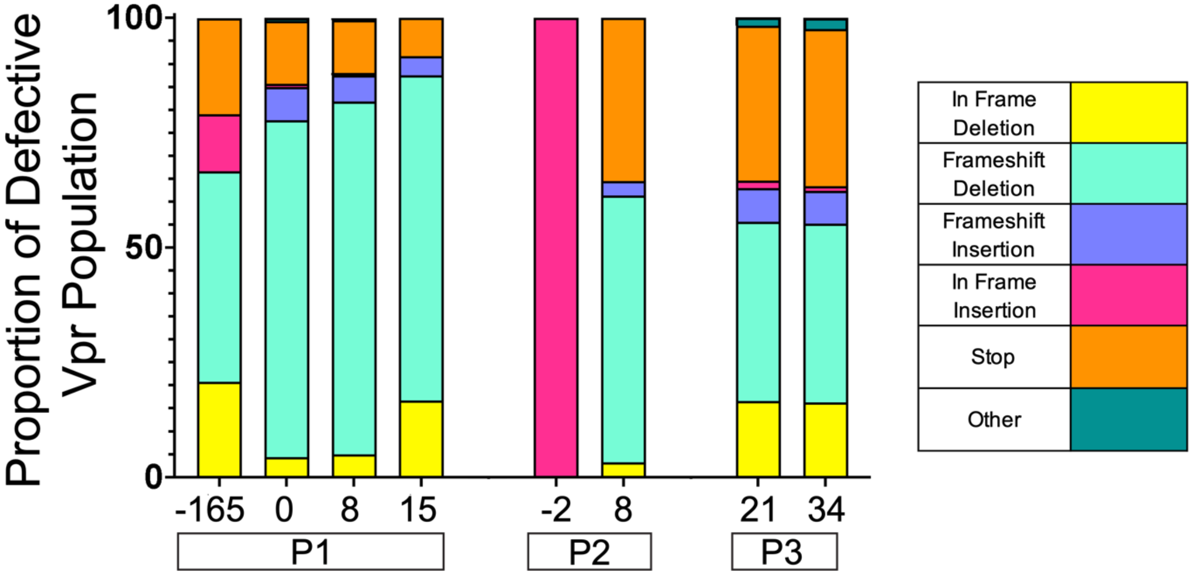


**Supplement Figure 4:** Proportion of *vpr* defects within the entire defective *vpr* gene population at each sampled timepoint. Proportions determined by Miseq Primer ID sequencing.
